# Detecting isoform-level allelic imbalance accounting for inferential uncertainty

**DOI:** 10.1101/2022.08.12.503785

**Authors:** Euphy Wu, Noor P. Singh, Kwangbom Choi, Mohsen Zakeri, Matthew Vincent, Gary A. Churchill, Cheryl L. Ackert-Bicknell, Rob Patro, Michael I. Love

## Abstract

Allelic imbalance (AI) of gene expression in heterozygous individuals is a hallmark of *cis*-genetic regulation, revealing mechanisms underlying the association of non-coding genetic variation with downstream traits, as in GWAS. Most methods for detecting AI from RNA-sequencing (RNA-seq) data examine allelic expression per exonic SNP, which may obscure imbalance in expression of individual isoforms. Detecting AI at the isoform level requires accounting for inferential uncertainty (IU) of expression estimates, caused by multi-mapping of RNA-seq reads to isoforms and alleles. *Swish*, a method developed previously for differential transcript expression accounting for IU, can be applied in a paired setting to detect AI. However, in AI analysis, most transcripts will have high IU across alleles such that even methods like *Swish* will lose power. Our proposed method, *SEESAW*, offers AI analysis at various level of resolution, including gene level, isoform level, and optionally aggregating isoforms within a gene based on their transcription start site (TSS). This TSS-based aggregation strategy strengthens the signal for transcripts that may have high IU with respect to allelic quantification. *SEESAW* is primarily designed for experiments with multiple replicates or conditions of organisms with the same genotype, as in an F1 cross or time course experiments of cell lines. Additionally, we introduce a new test for detecting AI that changes across a continuous covariate, as in a time course experiment. The *SEESAW* suite of methods is evaluated both on simulated data and applied to an RNA-seq dataset of differentiating F1 mouse osteoblasts.

## Introduction

Genome-wide association studies (GWAS) have identified tens of thousands of genomic loci that are associated with complex traits or diseases, many of which are located in non-coding regulatory regions [1]. One potential mechanism by which allelic variation in these non-coding regions may affect phenotype is that the variants reside in transcription factor (TF) binding sites and influence the activities of TFs and transcription.

Such a non-coding region may be referred to as a *cis*regulatory element (CRE). For individuals that are heterozygous at such a variant, the individual may exhibit imbalanced allelic expression at any genes regulated by the CRE. With RNA-sequencing (RNA-seq) experiments, it is possible to observe such imbalance in allelic expression in the sequenced reads if the same individual also is heterozygous for a variant in the exons of a regulated gene; other mechanisms of allelic imbalance, e.g. sequences affecting splicing or post-transcriptional regulation, are also possible to be detected. Recent advances in long-read technologies enable reconstruction of individual diploid genomes/transcriptomes, which leads to more accurate analysis at allele and isoform resolution. Analysis of allelic imbalance (AI) has the potential for higher power for detecting *cis*-genetic regulation regulation than analysis of total expression, as *trans*-regulatory and non-genetic effects on expression level are controlled for when comparing the two alleles within samples [2–8]. Such effects controlled for in AI analysis include both biological variability as well as technical artifacts that may distort total expression levels across genes. AI is also a powerful analysis to reveal *cis*-genetic regulation in heterozygous individuals that varies across samples representing different conditions, tissues, spatial contexts, or time periods [9–13].

AI can be isoform-specific, but due to low statistical power and challenges to statistical inference caused by multi-mapping, AI is often measured at the exon-level or gene-level. If different isoforms are subject to regulation from different sets of CRE, and these harbor genetic variants for which the individual is heterozygous, then such isoforms may exhibit different strength or direction of AI, as has been observed recently in an analysis of genomic imprinting at isoform level [14] and in a survey of expression in GTEx using long-read technology [15]. Being able to detect AI at the isoform level could provide insight into mechanisms of complex traits and disease. One challenge is that AI can only be observed when the individuals under study are heterozygous at an exonic variant. Furthermore, only a subset of reads that can be aligned or probabilistically assigned to a transcript or gene will provide allelic information. As described by Raghupathy et al. [16], an RNA-seq read can fall into various categories of multi-mapping with respect to gene, isoform, and allele, providing different information for expression estimation or “quantification” at different levels. The uncertainty in measuring the expression level from multi-mapping is referred to here as inferential uncertainty. When variants lie in exons that are not constitutive, reads overlapping these exonic variants provide information for isoform-level AI. In the other case, if exonic variants lie in constitutive exons, or in exons of dominant isoforms, AI can be effectively detected by existing methods such as *phASER* [8] or *WASP* [17], which count reads aligning to gene haplotypes. *WASP* for example examines imbalance in the pileup of reads mapped to the genome, after correction for technical bias due to differential mapping rates of the two alleles [18]. *WASP* can be followed up by methods such as *ASEP* for statistical inference of gene-level AI across a population of individuals that may be homozygous or heterozygous for regulatory variants [19].

A subset of existing methods are able to detect *cis*-genetic regulation at sub-gene resolution from short-read RNA-seq. Paired Replicate Analysis of Allelic Differential Splicing Events (*PAIRADISE*) can extract more information about allelic exon inclusion events, by counting reads that overlap both a exonic variant for which a subject is heterozygous, and an informative splice junction [20]. Within the *PAIRADISE* framework, reads are mapped to personalized genomes based on phased genotypes. *PAIRADISE* provides a statistical model for detecting allele-specific splicing events, by aggregating allelic exon inclusion within individuals, and builds upon their previous method GLiMMPS to detect splicing quantitative trait loci (QTL) across donors of all genotypes [21]. Two potential limitations for this approach are that such an approach is not able to aggregate allelic information along the length of a transcript or from the two paired ends of a fragment, and some cases of isoform-level AI may be missed when focusing on reads overlapping splice junctions, such as allele-specific differences in length of 5’ or 3’ untranslated regions (UTR).

Other method publications that have demonstrated quantification of expression at allelic- and isoform-level include *EMASE* [16], *kallisto* [22], *mmseq* [23], and *RPVG* [14]. *EMASE* proceeds in a similar manner to *PAIRADISE*, by first constructing a diploid reference, however in this case *EMASE* aligns reads to a diploid transcriptome, constructed via the *g2gtools* software. The *EMASE* authors found that hierarchical assignment of reads based on their information content in some cases outperformed equal apportionment as would occur using EM-based algorithms such as *RSEM* [24], *kallisto* [22], and *Salmon* [25] with a diploid reference transcriptome. *mmseq* allows for alignment of reads to a diploid reference transcriptome using *Bowtie* [26], and additionally can take into account gene-, isoform-, and allelic-multi-mapping when performing inference across alleles in its *mmdiff* step [27]. *mmdiff* computes posterior distributions of expression of each feature via Gibbs sampling. Features can be aggregated at various level of resolution by summing the posterior expression estimates within each sample. Aggregation also has proved an effective strategy in non-allelic contexts, as demonstrated in *tximport* [28], *SUPPA* [29], and *txrevise* [30]. *mmseq* also provides a method *mmcollapse* [27] to perform *data-driven* aggregation of features to reduce marginal posterior variance, although this procedure cannot currently be combined with differential analysis across alleles (i.e. AI analysis).

RNA isoforms may differ by transcription start sites (TSS), internal splice junctions, or termination sites. With the objective of detecting isoform-level AI, we introduce a strategy to group isoforms based on their transcription start sites (TSS). For a particular case of simulated AI, we show aggregating isoform-level expression estimates to the TSS offers better resolution than gene- or exon-level analysis, while exhibiting reduced inferential uncertainty compared to isoform-level analysis. We describe a suite of methods, Statistical Estimation of Allelic Expression using *Salmon* and *Swish* (*SEESAW*), for allelic quantification and inference of AI patterns across samples. *SEESAW* utilizes *Salmon* [25] to estimate expression with respect to an allele-specific reference transcriptome, and a non-parametric test *Swish* [31] to test for AI. *Swish* incorporates inferential uncertainty into differential testing and makes no assumption of the distributional model of the data. *SEESAW* follows the general framework of *mmseq* and *mmdiff* for haplotype- and isoform-specific quantification and uncertainty-aware inference. Here, the *SEESAW* methods were applied to simulation data to benchmark against previously developed methods for detection of AI within heterozygous individuals, making use of multiple individuals as biological replicates. We applied *SEESAW* to a mouse F1 cross time course dataset, where it detected genes containing both gene-level AI and isoform-level AI. *SEESAW* can detect cases of AI that are consistent across all samples, differential AI across two groups of samples, or dynamic AI over a covariate, with a new correlation-based test. *SEESAW* is made available within the *fishpond* package on Bioconductor, and is accompanied with a detailed software vignette on performing allelic analysis.

## Methods

### SEESAW

We first describe the steps in *SEESAW*, which combines both existing and new functionality across a number of software packages (Figure 1). *SEESAW* assumes that phased genotypes are available, and is primarily designed for multiple replicates or conditions of organisms with the same genotype. This can occur with multiple replicates of an F1 cross, or cell lines from individual human donors across developmental time points [32–34], or across conditions [35–38]. SEESAW facilitates importing the quantification data at various levels of aggregation: no aggregation (labelled here-after “isoform”, or equivalently “transcript”/”txp”), transcription start site aggregation (“TSS”), or gene-level aggregation (“gene”). The following steps were applied to analyze isoform-, TSS-, or gene-level AI. A sample-specific diploid transcriptome was constructed using *g2gtools* with the following input data: a reference genome (FASTA file); haplotype-specific variants (SNP and indel VCF files); and a catalog of all possible transcripts in the reference genome (GFF or GTF file). *g2gtools* was used to patch and transform the reference genome using the SNPs and indels from the the VCF file, and to extract transcripts from each haplotype of the custom diploid genome. Combined, these transcripts form the custom diploid transcriptome used to quantify RNA-seq reads. *Salmon* [25] was used to quantify expression at the level of allelic transcripts, where both alleles are kept in the reference during indexing (`--keepDuplicates`). This approach for allelic quantification, mapping reads to a custom diploid transcriptome, has been demonstrated as a successful strategy in previous work [23, 39, 22, 16], similarly for mapping reads to a spliced pangenome graph [14] or to a custom diploid genome for allelic read counting [40, 7, 41–43]. During the quantification step, 30 bootstrap replicates were generated to capture inferential uncertainty across genes, isoforms, and alleles (`--numBootstraps 30`).

**Figure 1.**
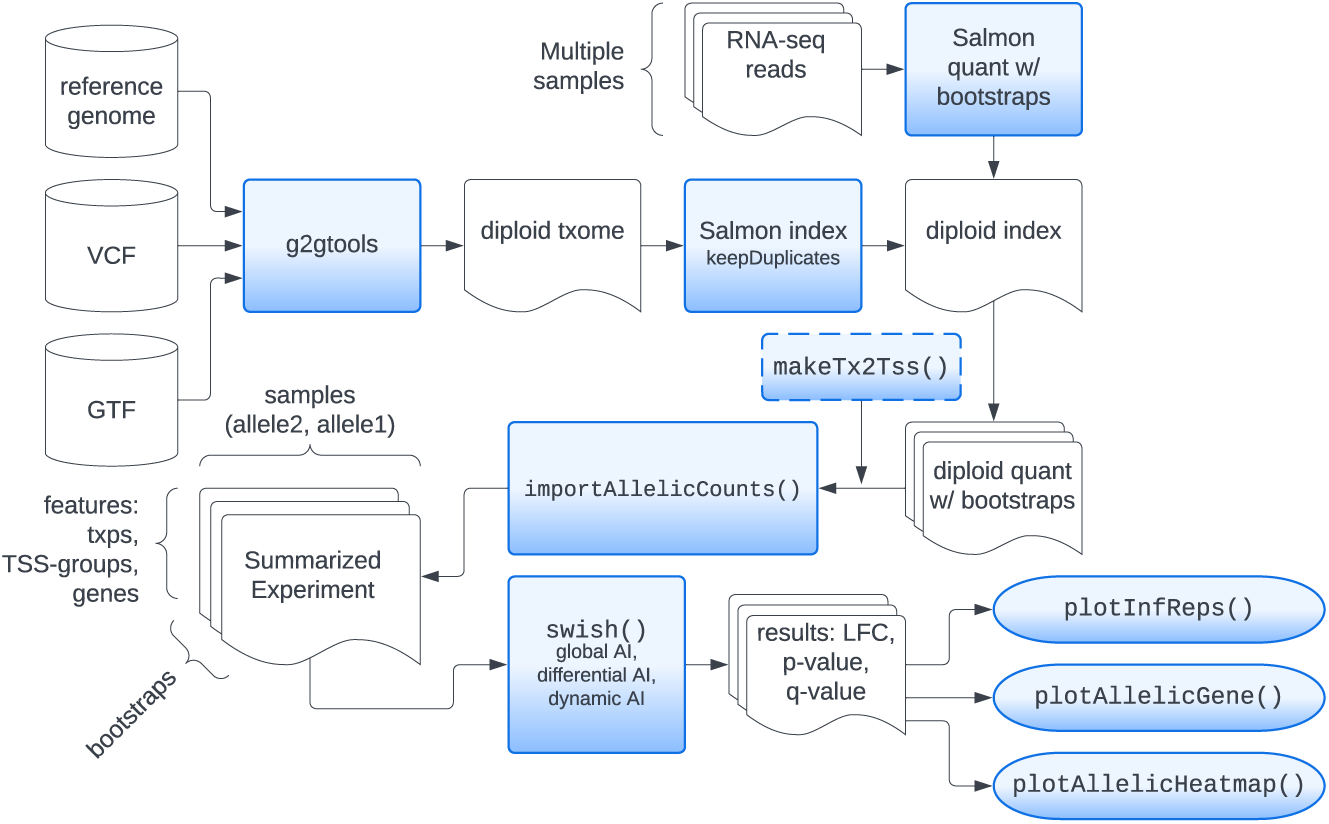
*SEESAW* pipeline for generating allelic expression estimates and performing statistical testing for allelic imbalance.

To increase statistical power in testing for AI at sub-gene resolution, we recommend to group isoforms at a resolution that prioritizes discovery of *cis*-genetic regulation effects from non-coding variation in the promoter or in CRE that affect a particular promoter. Aggregation of isoform expression to higher levels has been shown to reduce inferential uncertainty and may improve detection power as long as the signal of interest is not at the same time lost or diminished through aggregation [27, 28, 44]. Other related approaches include performing inference at the level of equivalence classes (partitions of reads) [45, 46], although here we focus on analysis performed over sets of one or more transcripts for their biological interpretability. During aggregation, both counts and bootstrap replicate counts were summed across isoforms within a TSS-based group for each allele. TSS groups can be defined strictly (identical start position) or with some basepair (bp) tolerance (“fuzzy TSS groups”). After aggregation, every aggregate feature will have a point estimate of abundance as well as a vector representing the bootstrap distribution for each of the two alleles. Likewise, we explored gene-level aggregation, summing across all isoforms for a gene.

In the *fishpond* Bioconductor package, we used a convenience function `makeTx2Tss` for generating TSS groups (with an optional parameter to group nearby TSS), and then used `importAllelicCounts` to import the estimated counts, abundances, and bootstrap replicates, producing a *SummarizedExperiment* object and leveraging the *tximeta* package [47, 48]. In the case that there was no read information to distinguish the two alleles, e.g. identical sequence, or no reads covering any sequence differences, *Salmon* splits the total counts equally among the two alleles, so the estimated allelic fold change was equal to 1. Such features were filtered out of the dataset before differential testing, as demonstrated in the software vignette. Prior to differential testing, features that did not have a minimum count of 10 for three or more samples were filtered out.

*Swish* [31] was used here to detect AI across biological replicates or conditions while taking into account inferential uncertainty. *Swish* is a nonparametric method originally designed for isoform-level differential expression that extends the gene-level method *SAMseq* [24]. We tested for the existence of AI (allelic fold change not equal to 1) for a given feature across all samples by specifying a paired analysis with `x=“allele”, pair=“sample”`, which was referred to as “Global AItesting”. Reviewing the paired-sample method developed in Zhu et al. [31], *Swish* for global AI used the Wilcoxon signed-rank test [49], where the paired data in this context are the allelic counts for each sample and each bootstrap. The signed-rank test statistic was averaged over bootstraps (in this case, 30 bootstraps), and the allelic labels were swapped to generate a permutation null distribution. The framework of averaging nonparametric test statistics over replicate datasets, followed by permutation-based *q*-value computation was derived from *SAMseq* [50].

We further extended *Swish* to test for changes in AI along a continuous covariate. We tested for non-zero correlation between the log allelic fold change within paired samples and a continuous covariate by specifying e.g., `x=“allele”, pair=“sample”, cov=“day”`, which was referred to as “Dynamic AI testing”. Either Pearson or Spearman correlations can be computed between the pairwise log fold changes and the continuous covariate, and this statistic was then averaged over bootstrap samples. To stabilize the log fold change, a pseudocount was added to the numerator and denominator of the allelic fold change, with a default value of 5, as has been used previously for bulk RNA-seq [31]. The continuous covariate was then permuted and correlations recalculated to generate a null distribution over all permutations and all features, followed by *q*-value computation with the *qvalue* package [51] to obtain the significance of the relationship between the allelic fold change with the continuous covariate. Testing for changes in AI across a categorical covariate was already available in *Swish* as an “interaction test” and is demonstrated in the software vignette on allelic analysis.

For statistical methods designed for comparing gene expression across samples, it is common practice that measures of gene expression are scaled or an offset is included in the model to account for a well-known technical bias: differences in sequencing depth across samples affect the observed or estimated counts and if left unadjusted the across-condition estimates would be biased. Since *SEESAW* focuses on testing differences in expression between the two alleles within the same sample, the sequencing depth bias affected both allelic counts equally, and scaling/offsets were not needed. Thus, this scaling step should not be performed.

A number of plotting functions in *fishpond* were used to facilitate visualization of allelic expression changes across samples, isoforms, and covariates. `plotInfReps` was used to visualize allelic expression estimates and inferential uncertainty across samples and conditions/time points [31]. `plotAllelicGene` was used to visualize isoform- or TSS-group-level allelic expression data along with a diagram of a given gene model, using the *Gviz* package [52]. `plotAllelicHeatmap`, was used to visualize allelic expression across isoforms or TSS groups and samples, leveraging the *pheatmap* package [53]. In *SEESAW*, transcript ranges were represented using `GRanges` [54] objects generated from `TxDb` or `EnsDb` databases [55], and attached as `rowRanges` to the main dataset object with estimated counts, abundance and bootstrap counts, facilitating downstream plotting and data exploration.

### Simulation

To assess the performance of different methods in recovering gene-level and isoform-level AI, we simulated RNA-seq reads from a diploid transcriptome derived from the *Drosophila melanogaster* reference transcriptome, restricted to chromosomes 2, 3, 4, and X, simulating RNA transcripts from female flies. The simulation contained a total of 10 samples, with an average sequencing depth of 50 million paired-end reads per sample. The maternal reference transcripts included the protein-coding and non-coding RNA from Ensembl [56] (release 100), and the paternal reference transcripts were created from maternal transcripts by adding single nucleotide variants. To create paternal alleles, we randomly selected 5 exons from each gene, and the mid-position nucleotide of the selected exons was substituted with its complement nucleotide. Genes overlapping simple tandem repeats of size 50 bp or larger were excluded from the simulation, leaving 14,821 genes.

While the majority of genes (93.3%) were simulated to not have AI, two types of AI were simulated: “concordant AI”, where all isoforms had concordant allelic fold change, and “discordant AI”, where there were discordant allelic fold changes among the isoforms of the gene. In the discordant AI case, the RNA abundance was balanced across the two alleles when summed across all the isoforms of the gene. We randomly selected 1,000 genes using the following criterion: 1) the number of annotated isoforms in the selected gene was between three and six, 2) the gene had at least two isoforms sharing the same transcription start site (TSS), 3) the gene had at least two distinctive TSS. Half of the selected genes (500) were simulated to have concordant AI and the other half (500) were simulated to have discordant AI, and the remaining 13,821 genes were simulated to have allelic balance.

The abundance of the maternal allele was set to a constant value, and the paternal allelic abundance was altered to generate AI. For genes with concordant AI, the paternal allele was either 25% up-regulated or down-regulated, chosen at random per gene. Within each gene with discordant isoform-level AI, one TSS was randomly chosen and isoforms sharing the selected TSS had abundance increased on the paternal allele. Abundance was increased such that the up-regulation fold change was equal to 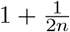, where *n* is the number of isoforms with the selected TSS. The other isoforms of the gene had paternal abundance decreased at an equal rate such that the gene-level abundance was kept constant. Expected count values were then generated from the alleles of all transcripts by multiplying abundance by the transcript length and scaling up to the desired library size. Reads were generated using *polyester* [57], with the following settings: fragments with a mean size of 400 bp, paired-end reads of 150 bp, and Negative Binomial dispersion parameter `size=100`, such that the simulation contained across-sample biological variation on the allelic counts and the allelic fold change. Paired-end reads were shuffled so that the reads were listed in a random order. Read generation and all subsequent analysis steps for *SEESAW* and other methods were automated using a *Snakemake* workflow [58] available at github.com/mikelove/ase-sim.

We applied the *SEESAW* pipeline as well as existing methods *mmseq* and *WASP* to the simulated data. As our interest in this work was in identifying isoforms affected by *cis*-genetic regulation, we focused our comparisons on methods that aim to aggregate allelic information along the entire extent of the transcribed region, whereas other existing methods have focused on allelic differences in internal splice events (splicing quantitative trait loci, or sQTL); we chose a method that attempts to resolve AI at the isoform level (*mmseq*), as well as a method that resolves bias from genomic multi-mapping reads (*WASP*) and is primarily focused on gene-level AI. *SEESAW* and other methods were provided with the complete exonic sequence of the two alleles, and the gene annotation, either as FASTA (for *Salmon* and *mmseq*) or VCF files with known phasing (for *WASP*). We also utilized the ground truth of simulation to obtain an optimal isoform-grouping strategy, called “oracle”. For “oracle” grouping, up-regulated or down-regulated isoforms within a gene were grouped in cases of discordant AI. Otherwise, all isoforms within a gene were grouped together.

The following steps were used to apply *mmseq* (version 1.0.10a) and its differential testing step *mmdiff* to the simulated data. *Bowtie* (version 1.3.1) [26] was used to index the diploid reference transcripts and to align the reads. During the alignment, only the alignments that fell into the best stratum were reported if the alignments fell into multiple stratum using `–best --strata`. If more than 100 reportable alignments existed for a particular read, then all alignments were suppressed using the `-m` option. After obtaining *mmseq* expression estimates at gene and isoform level, we manually separated the estimates for the maternal and paternal alleles, and subsequently used `mmdiff` to test for differential expression between the two alleles, using the flags `-de 10 10 <maternal files> <paternal files>`. Posterior probability of equal expression was used to threshold and define significant sets of transcripts or genes.

We ran *WASP* on the simulated data according to its recommended usage. First, HISAT2 (version 2.2.1) [59] was used to align reads to the bdgp6 reference genome, downloaded from the HISAT2 website. An h5 database was created from the simulation VCF file containing the location of the exonic SNPs and the known phasing information. HISAT2 was used to re-align the reads with flipped nucleotides, and genomic multi-mapping reads that would otherwise bias allelic ratios were filtered. Read counts and heterozygous probabilities were adjusted using *WASP* scripts. Finally the Combined Haplotype Test (CHT) [17] was applied with recommended defaults `--min_counts 50` and `--min_as_counts 10` to generate a *p*-value per gene for AI across samples. Multiple testing was controlled via the *locfdr* package [60], applied to z-scores derived from *WASP p*-value output.

To visually compare the AI simulation results across methods, we used the iCOBRA [61] Bioconductor package. We assessed the performance according to the true and reported allelic status of the transcripts (balanced or imbalanced), where reported significance of AI of a gene or TSS group was propagated to its isoforms. As the simulation consisted only of genes in which all isoforms or no isoforms exhibited true AI, this approach to compare methods at the transcript level should not unfairly impact the performance of the aggregated (gene- or TSS-level) AI tests.

### Osteoblast differentiation time course

We applied *SEESAW* to an RNA-seq dataset of primary mouse osteoblasts undergoing differentiation, from F1 C57BL/6J x CAST/EiJ mice, to assess both global and dynamic AI. In the differentiation experiment, which has been described previously [62, 63], pre-osteoblast-like cells were extracted from neonatal calvaria, and cells were FACS sorted based on expression of CFP, which was under the control of the Col3.6 promoter. Differentiation was induced with an osteoblast differentiation cocktail in sorted cells and RNA was collected every two days from day 2 to day 18 post differentiation (nine time points). Three technical replicates per time point were combined and quantified together as one biological replicate, after quality checking with *FASTQC* and *MultiQC* [64]. Expression data for osteoblasts from C57BL/6J mice of the same experiment are publicly available on the Gene Expression Omnibus (GSE54461).

Reference transcripts were generated via *g2gtools* using a reference genome, strain-specific VCF files, and the reference gene annotation. The GRCm38 primary assembly for *Mus musculus* was downloaded from Ensembl (release 102) [56], and strain-specific VCF files `CAST_EiJ.mgp.v5.snps.dbSNP142.vcf` (SNP) and `CAST_EiJ.mgp.v5.indels.dbSNP142.normed.vcf` (indel) for mm10 were downloaded from the Mouse Genomes Project [65] ftp://ftp-mouse.sanger.ac.uk/REL-1505-SNPs_-Indels/. Reference transcripts (`Mus_-musculus.GRCm38.102.gtf`) were downloaded from Ensembl (release 102) [56], and subsequently were patched and transformed for the CAST/EiJ strain using *g2gtools* (version 0.2.7). All code including a *Snakemake* workflow [58] for generation of the diploid transcriptome is provided in the `diploid_txomes` directory at github.com/mikelove/osteoblast-quant. *Salmon* was used to quantify the RNA-seq reads against the custom diploid transcriptome with 30 bootstrap replicates, and these data were imported into Bioconductor and analyzed with *Swish* as described in the *SEESAW* section above. *Swish* with global AI test was performed on isoform level, TSS-group level and gene level, and results were compared at various levels of resolution. In addition, we used the newly developed feature in *Swish* to test for dynamic AI: we tested the correlation between the log fold change comparing across alleles within a sample and the day of differentiation, using the Pearson correlation.

## Results

### Simulation

Simulation of an F1 cross based on the *Drosophila melanogaster* transcriptome was used to assess method performance when the true AI status of each transcript was known. iCOBRA diagrams [66] were used to assess the sensitivity, or true positive rate (TPR), and the false discovery rate (FDR) at nominal FDR thresholds of 1%, 5%, and 10%. Sensitivity was assessed per transcript, where detection of AI for a gene-level method was propagated to each of the gene’s expressed isoforms. We used Integrative Genomics Viewer (IGV) [67] to visualize the distribution of HISAT2 [59] aligned reads along the reference genome, after removing allelic-biased multi-mapping reads with *WASP* [17]. While *SEESAW* uses reads mapped to the diploid transcriptome with *Salmon*, examining genome-aligned reads with IGV allowed us to identify examples of reads that contained both allelic- and isoform-level information (Figure S1).

As expected, *SEESAW* had the strongest power to detect AI when oracle information about the grouping of transcripts by AI signal was used to aggregate allelic signal (“oracle” in Figure 2A). Among methods that were not provided information about the true grouping of transcripts by AI signal, *SEESAW* with TSS aggregation or gene-level aggregation, gene-level *mmdiff*, and *WASP* had similar sensitivity at 1% nominal FDR, and these methods had observed FDR for this nominal cutoff in the range of 0-2%. Notably, *SEESAW* with TSS aggregation had the highest overall sensitivity at 5% and 10% nominal FDR, above any of the gene- or isoform-level methods. The reason behind the higher overall sensitivity can be seen when stratifying by types of AI, as in Figure 2B; *SEESAW* with TSS aggregation was able to detect discordant AI on isoforms within a gene that could be masked after aggregation to the gene level. Gene-level *SEESAW*, gene-level *mmdiff*, and *WASP* had loss of sensitivity to detect these discordant cases of AI. However, these three methods had higher sensitivity than *SEESAW* using TSS aggregation when AI was concordant across all isoforms of a gene. This is expected as aggregation at the appropriate level strengthens the AI signal while reducing inferential uncertainty, so increasing power. Both *SEESAW* and *mmdiff* at the isoform level did not have as high sensitivity as methods that aggregated signal. UpSet diagrams [68] of the sets of transcripts called by each method compared to the true AI transcripts indicated the highest overlap among the gene-level methods and TSS or oracle aggregation (Figure S2).

**Figure 2.**
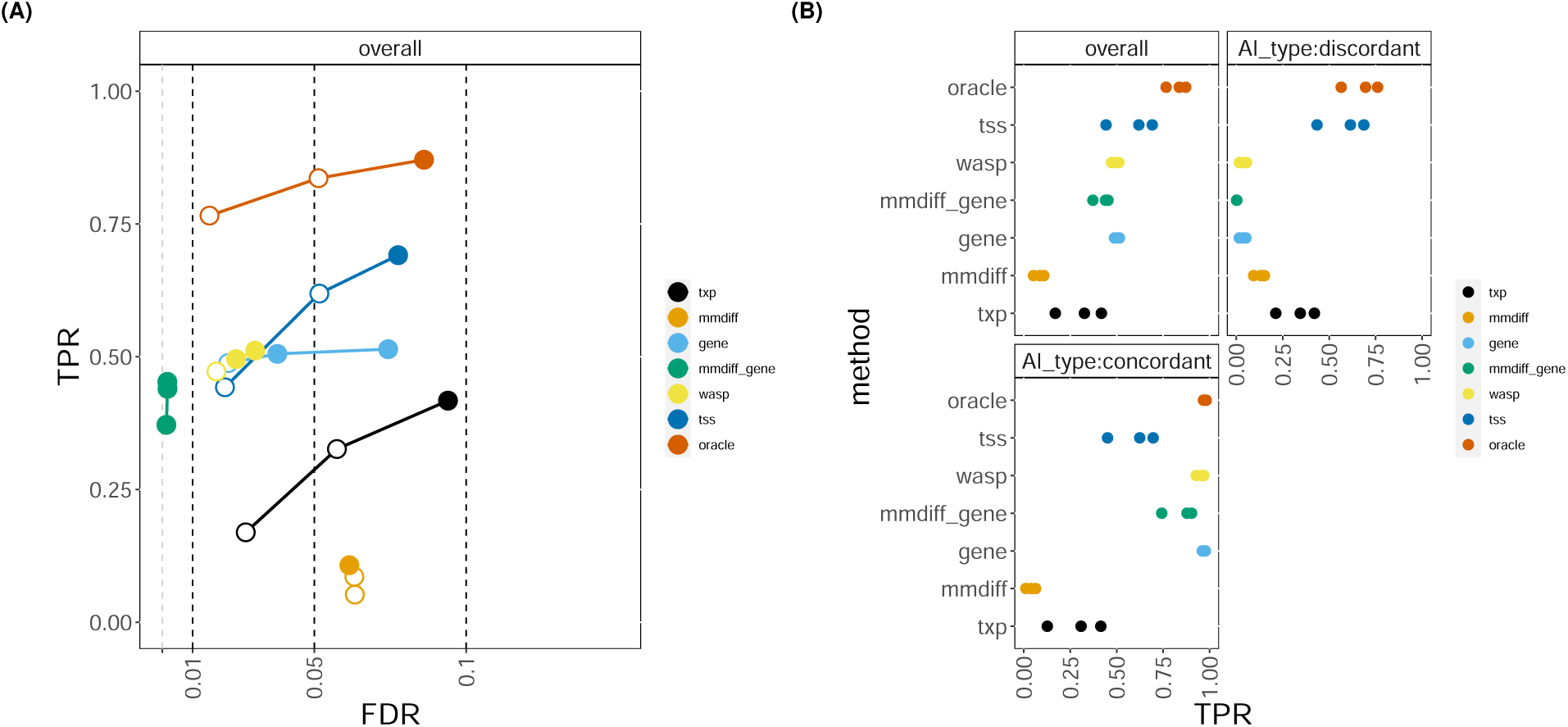
Comparing results of *SEESAW* on *polyester* simulation to *mmdiff* and *WASP*. A) iCOBRA plot of sensitivity (true positive rate, or TPR) over achieved false discovery rate (FDR) with three circles indicating 1%, 5%, and 10% nominal FDR cutoffs, respectively. Filled circles indicate observed FDR less than nominal FDR. B) Overall sensitivity for all cases of AI and sensitivity stratified by type of AI: “discordant” AI across isoforms within a gene or “concordant” AI within gene.

We additionally assessed performance of *SEESAW* compared to a new inference pipeline from the *WASP* developers, called *WASP2* (Figure S3). *WASP2* was equally sensitive as *WASP* in detecting gene-level AI, while it had less sensitivity than *SEESAW* to detect discordant AI signal as expected since it follows a similar approach to *WASP*. While we used the *locfdr* package [60] for multiple test correction for *WASP*, we found Benjamini-Hochberg [69] correction performed well for computing FDR-bounded sets for *WASP2*.

### Osteoblast differentiation time course

Following creation of the diploid reference and quantification steps of the *SEESAW* pipeline, we used *Swish* to test for consistent AI across all time points of the osteoblast differentiation dataset. While exploring the osteoblast differentiation data, we observed that for isoforms of a gene with TSS that were near each other (within 50 bp), these isoforms often shared similar estimated allelic fold change as calculated with *SEESAW*. To facilitate data visualization, strengthen biological signal, and reduce inferential uncertainty further, we chose to group any transcripts with TSS within 50 bp (“fuzzy TSS groups”). We tested at different levels of resolution: gene level, isoform level, and TSS level. To compare across these levels, we looked at genes in common: a gene was considered significant for global AI at isoform level or TSS group level if at least one isoform or TSS group within the gene was significant (nominal FDR < 5%). Isoform-level testing for global AI returned the most genes, with 6,116 significant genes, followed by gene-level with 5,701 genes, and TSS-level grouping with 5,573 genes. The majority of genes (4,625) were in common across all three levels of resolution (UpSet plot [68] provided in Figure S4).

Gene-level aggregation had high overlap with TSS-level indicating that, at least for global AI testing, most of the AI signal was not masked by discordant direction of AI among isoforms within a gene. Among genes displaying global AI under aggregation to the gene level, the TSS groups within those genes often had estimated imbalance in the same direction as the gene imbalance – 97.3% of significant genes had all of their TSS groups with significant AI having the same direction as the gene-level estimate. However, *SEESAW* was able to detect – among the 2.7% remaining genes – interesting examples of genes that had different direction of AI among its isoforms. A complete list of the 134 genes showing these significant and discordant patterns within gene is provided in Table S1 and in the Zenodo deposition. For example, *Fuca2* exhibited discordant AI with the CAST/EiJ allele more highly expressed than C57BL/6J for one of the two leftmost (more 5’) TSS but less expressed than C57BL/6J for the rightmost (more 3’) TSS, with both TSS groups significant at 5% FDR (Figure 3A).

**Figure 3.**
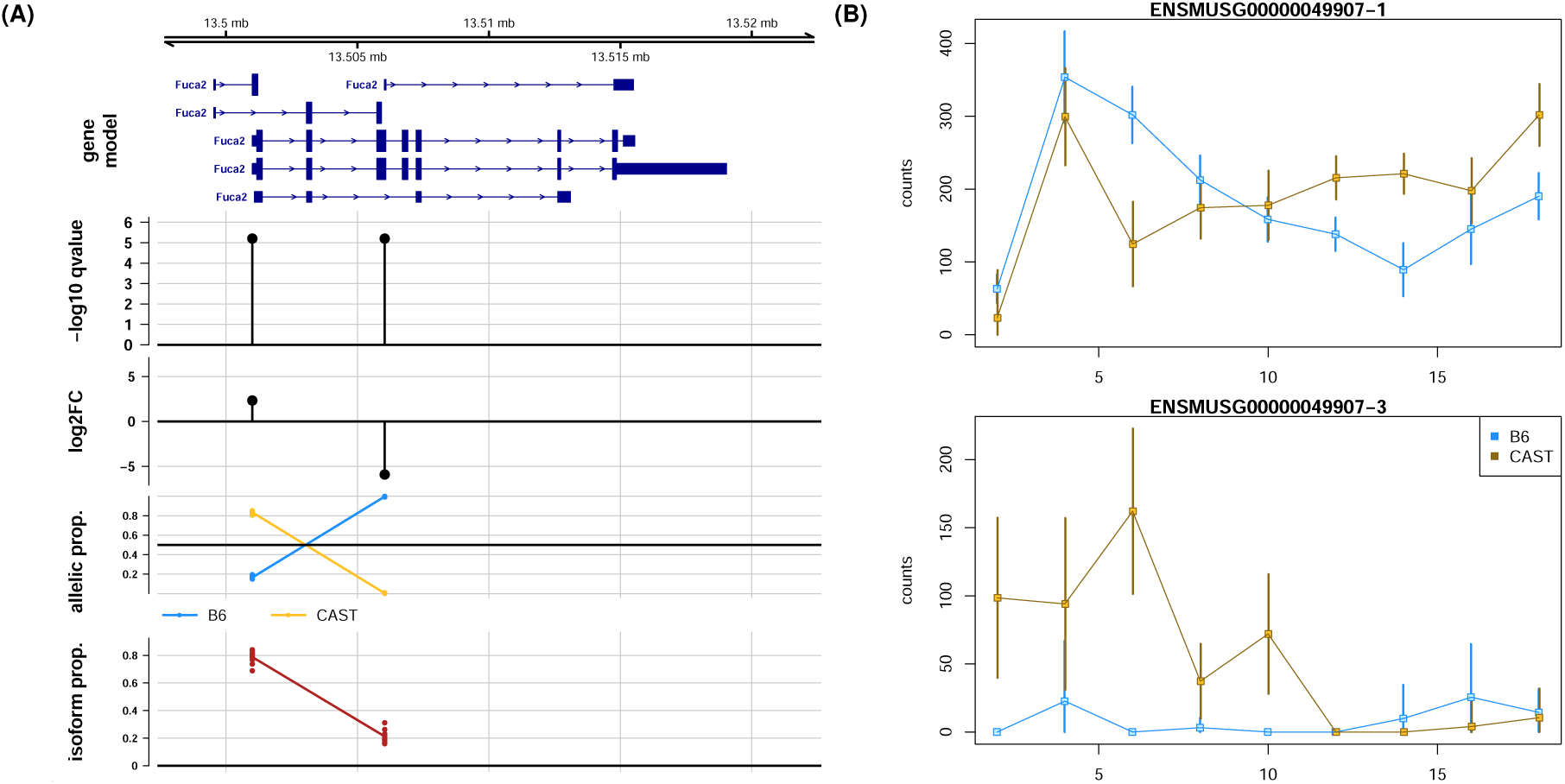
Application of *SEESAW* to osteoblast differentiation data at TSS level. A) Global AI results for *Fuca2* where isoform groups had discordant direction of imbalance. The computed statistics are plotted directly below the TSS group. Isoform proportion per TSS group was calculated from TPM (transcript per million). B) Isoform-level dynamic AI revealed for two TSS groups of *Rasl11b*. Estimation uncertainty shown with error bars (95% intervals based on bootstrap variance).

We additionally tested for dynamic AI using the correlation test implemented in *Swish*, again at gene level, isoform level, and TSS level. Gene-level dynamic AI testing returned the largest number of significant genes (nominal FDR <5%): 57 genes displayed dynamic AI at gene level, 49 genes at TSS level, and 23 at isoform level (Figure S5). Those significant genes shared across all levels only represented a third of those detected at gene level, where another third were shared only between gene level and TSS level. Thus TSS-level aggregation appeared to help recover signal that would be lost if only testing at the isoform level.

Interestingly, we detected genes such as *Rasl11b* that had isoform-level AI trending in different directions over time (Figure 3B, Figures S6 and S7). *Rasl11b* exhibited dynamic AI for two TSS groups, with the CAST/EiJ allele more lowly expressed than the C57BL/6J allele for TSS group “1” from day 2 to day 6, roughly balanced from day 8 to day 10, and finally with CAST/EiJ more highly expressed from day 12 to day 18. The other TSS group “3” had almost the opposite allelic ratio behavior: CAST/EiJ more highly expressed earlier in time but both alleles tending toward balanced, low expression at the end of the time course. While for *Rasl11b* this pattern was also significant when testing at the isoform level, other genes such as *Calcoco1* demonstrated the advantage of grouping features: *Calcoco1* exhibited dynamic AI for two TSS groups, “5” and “6”, which differed in the direction of change in the imbalance (Figures S8 to S10). Here, the *p*-value and *q*-value for TSS group “6” was reduced when aggregating counts from the isoform to TSS-group level.

## Discussion

Here we introduce a new suite of methods, *SEESAW*, to obtain allele-specific abundance with bootstrap replicates used to capture inferential uncertainty across genes, isoforms, and alleles, and to perform statistical testing of global or dynamic AI. We propose to aggregate estimates of allelic expression of isoforms by their TSS to increase statistical power in testing for AI that is a result of *cis*-genetic regulation on or within the promoter. We introduced two different AI testing procedures: global AI to test for the existence of consistent allelic fold changes across samples, and dynamic AI to test for non-zero correlation between the log allelic fold change and a continuous covariate. *SEESAW* can also be used to test differential AI between two groups, as introduced in Zhu et al. [31] and shown in the software vignette. The above tests utilize nonparametric testing that make no assumption on the distribution of the data itself, which had better performance than a standard beta-binomial generalized linear model (data not shown). In simulation, we demonstrated that *SEESAW* on TSS level had the highest sensitivity in the case that AI was discordant within gene, and achieved an FDR that was close to the nominal value at all levels of resolution (gene-, TSS-, or isoform-level testing), implying *SEESAW* can maintain error control despite high and heterogeneous levels of uncertainty. *SEESAW* at gene level performed comparably to existing methods such as *WASP* and *mmdiff* at gene level. For the osteoblast differentiation experiment, *SEESAW* was able to recover some genes with discordant isoform-level AI across all time points, and was able to detect genes with isoform-level AI that changed over time in different directions. Currently, *SEESAW* does not support alignment of haplotypes across individuals of different genotype. SNP-based analysis simplifies this problem, but at a loss of information, as evidence of AI may be distributed across multiple exonic variants within a transcript. A newly developed approach *RPVG* [14] maps RNA-seq reads to a spliced pangenome, and then provides haplotype-specific transcript abundance estimates for each individual. It would require further work for the methods presented here to group individuals by their haplotype combinations per gene and perform across-sample inference while accounting for estimation uncertainty using *Swish*. Another limitation of our current approach is that grouping isoforms together based on their TSS reveals shared promoter-based regulation, but may miss isoform-specific AI caused by intronic variation or variation that affects nonsense-mediated-decay. *PAIRADISE* uniquely targets these cases of AI on splicing events, and could be considered for these cases. Alternatively, the framework of *SEESAW* can be adapted and used with other aggregation rules for different biological purposes, e.g. aggregating isoforms by various splicing events in a manner similar to *SUPPA* or *txrevise*. While *SEESAW* can be used at various levels of resolution, from transcript or TSS group up to gene level, if the focus of interest is gene-level AI, we found that *WASP* and *WASP2* were equally sensitive and had good control of false discoveries, using *locfdr* and Benjamini-Hochberg correction, respectively. Additionally, the *ASEP* [19] method can be run on allelic counts from *WASP*, and allows for detection of gene-level AI across a population using a mixture model to account for the unobserved regulatory variants – individuals that are heterozygous for exonic variants may be either homozygous or heterozygous for regulatory variants. The analytical consequences of multiple regulatory SNPs and varying degree of linkage disequilibrium (LD) of these to the exonic SNP, with respect to detection of AI, has been described previously by Xiao and Scott [4].

While here we relied on gene annotation to group together transcripts and reduce inferential uncertainty of allelic expression estimates, another approach would be to use data-driven aggregation methods such as *mmcollapse* and *Terminus* [44]. We were not able to perform differential testing across alleles with *mmdiff* after aggregation with *mmcollapse*. A future direction that may improve performance with the inclusion of *Terminus* in the *SEESAW* pipeline would be stratification of different null distributions for test statistics in *Swish* based on aggregation level (transcript, transcript-group, or gene level).

After having detected AI, a natural next step is to try to understand the mechanism of *cis*-genetic regulation. It is possible to associate the AI seen on transcripts or genes with one or more regulatory variants, either through phasing or usage of population-level LD to establish the search space. The list of candidate regulatory SNPs can be further refined by integrating allelic signal at the epigenomic level, including allelic binding of proteins [40, 70–72], allelic accessibility [73, 74] or allelic methylation [75]. Alternatively, search for altered transcription factor binding motifs can be combined with RNA or protein abundance of potential regulators to winnow down the list of candidate causal regulatory variants [11, 76, 77]. It may also be of interest to detect in which cell types the allelic signal may be strongest or exclusively present, as has been investigated in recent methods for single cell allelic expression or accessibility datasets [78–80]. Finally, we note that a number of methods have shown that AI can be effectively integrated with total expression across individuals of all three genotypes [5, 81–84, 13]. This approach uses more information, and so should produce a gain in sensitivity, as well as extending beyond genes harboring exonic variants, which is a limitation for AI-based methods.

## Data Availability

The B6xCAST mouse osteoblast RNA-seq experiments have been submitted to SRA with the following BioSample accessions: SAMN29983440, SAMN29983441, SAMN29983442, SAMN29983443, SAMN29983444, SAMN29983445, SAMN29983446, SAMN29983447, SAMN29983448.

For the mouse osteoblast dataset, R data objects containing the total and allelic counts at gene and isoform level, diploid transcriptome sequences, *Salmon* quantification directories for all samples, global and dynamic AI test results at all three levels of resolution, and the discordant global AI gene list are provided at the following Zenodo accession number, doi: 10.5281/zenodo.6963809.

For the *Drosophila melanogaster* simulated samples, R data objects containing the simulated abundances and status of each transcript, simulated transcriptome sequences, *Salmon* quantification directories for all samples, SummarizedExperiment objects at various levels of resolution, and results tables for each method are provided at the following Zenodo accession link, doi: 10.5281/zenodo.6967130.

## Code Availability

Code for quantifying the osteoblast data is provided at https://github.com/mikelove/osteoblast-quant and code for AI testing and compiling results across different levels of resolution is provided at https://github.com/FennecFish/osteoblast-test.

Code for generating the simulated data is provided at https://github.com/mikelove/ase-sim and code for running *Swish* and compiling the results from other methods is provided at https://github.com/mikelove/swish-ase-assessment.

## Acknowledgments

We thank Dr. Matthew Hibbs for his contributions to the generation of the original dataset, and Dr. Anuj Srivastava and Sai Lek for their assistance with data deposition. We thank the following individuals for feedback and suggestions during the development of the method and manuscript: Dr. Hirak Sarkar, Dr. Andrew Parker Morgan, Dr. Fernando Pardo-Manuel de Villena, Dr. Terry Furey, Dr. Nil Aygun, Dr. Jason Stein, Dr. Kevin Currin, Dr. Karen Mohlke, Dr. Ina Hoeschele, Dr. Ernest Turro, Mr. Aaron Ho, Dr. Graham McVicker, and Dr. Bryce van de Geijn.

## Funding

NIH R01 HG009937, NSF CCF-1750472 and CNS-1763680

- Rob Patro
- Noor P Singh
- Mohsen Zakeri
- Michael I Love

NIH T32 CA106209

- Euphy Wu

NIH R01 GM070683

- Gary A Churchill
- Matthew Vincent

## Supplementary Information

**Figure S1.**
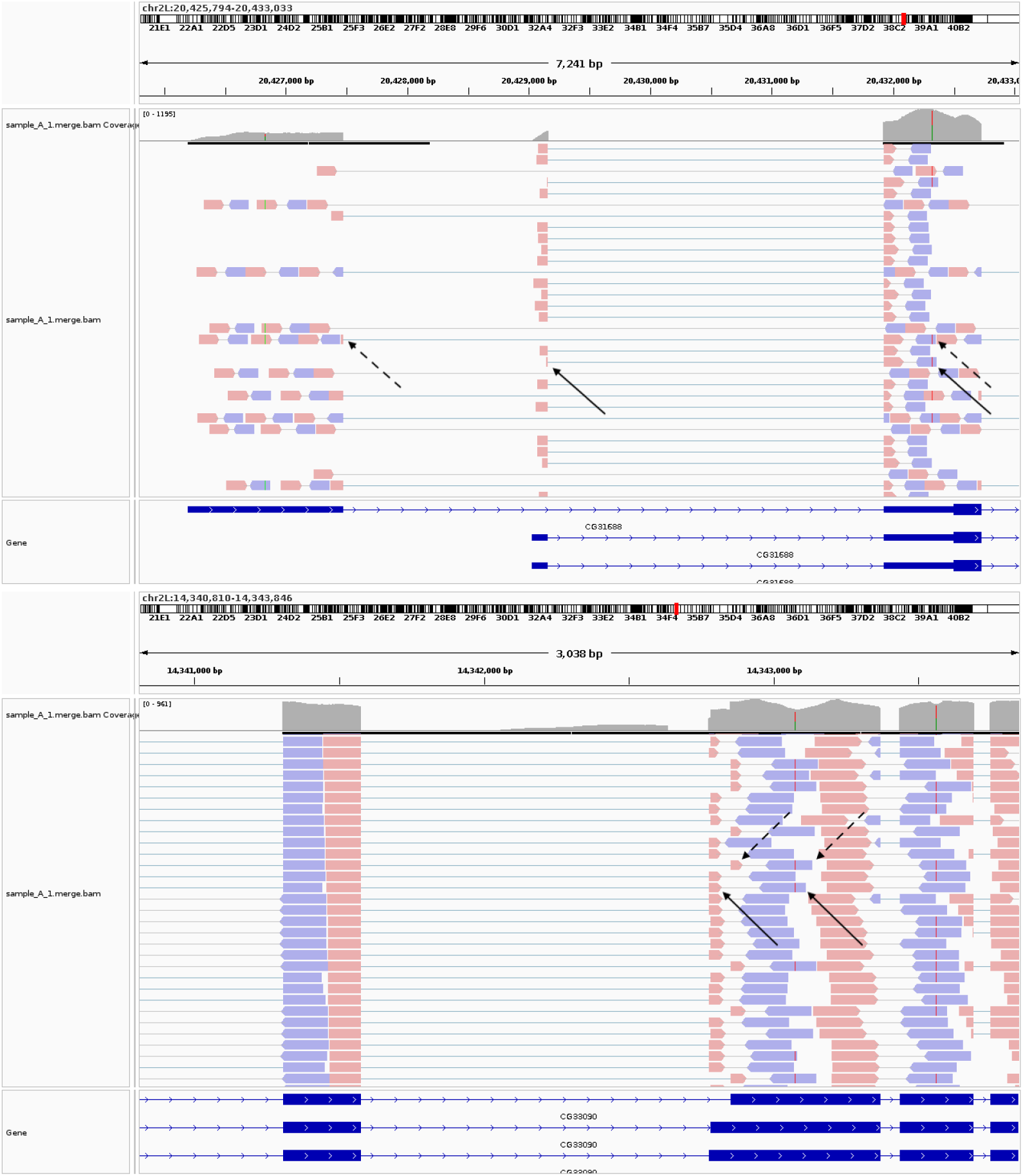
IGV visualization of HISAT2 aligned reads for the simulated *Drosophila melanogaster* dataset. The two depicted loci are *CG31688 / FBgn0263355* (top) and *CG33090 / FBgn0028916* (bottom). Paired-end reads such as those highlighted with dashed and solid arrows provide information to both isoform and allelic expression, used by *Salmon* to distribute isoform- and allelic-multi-mapping reads.

**Figure S2.**
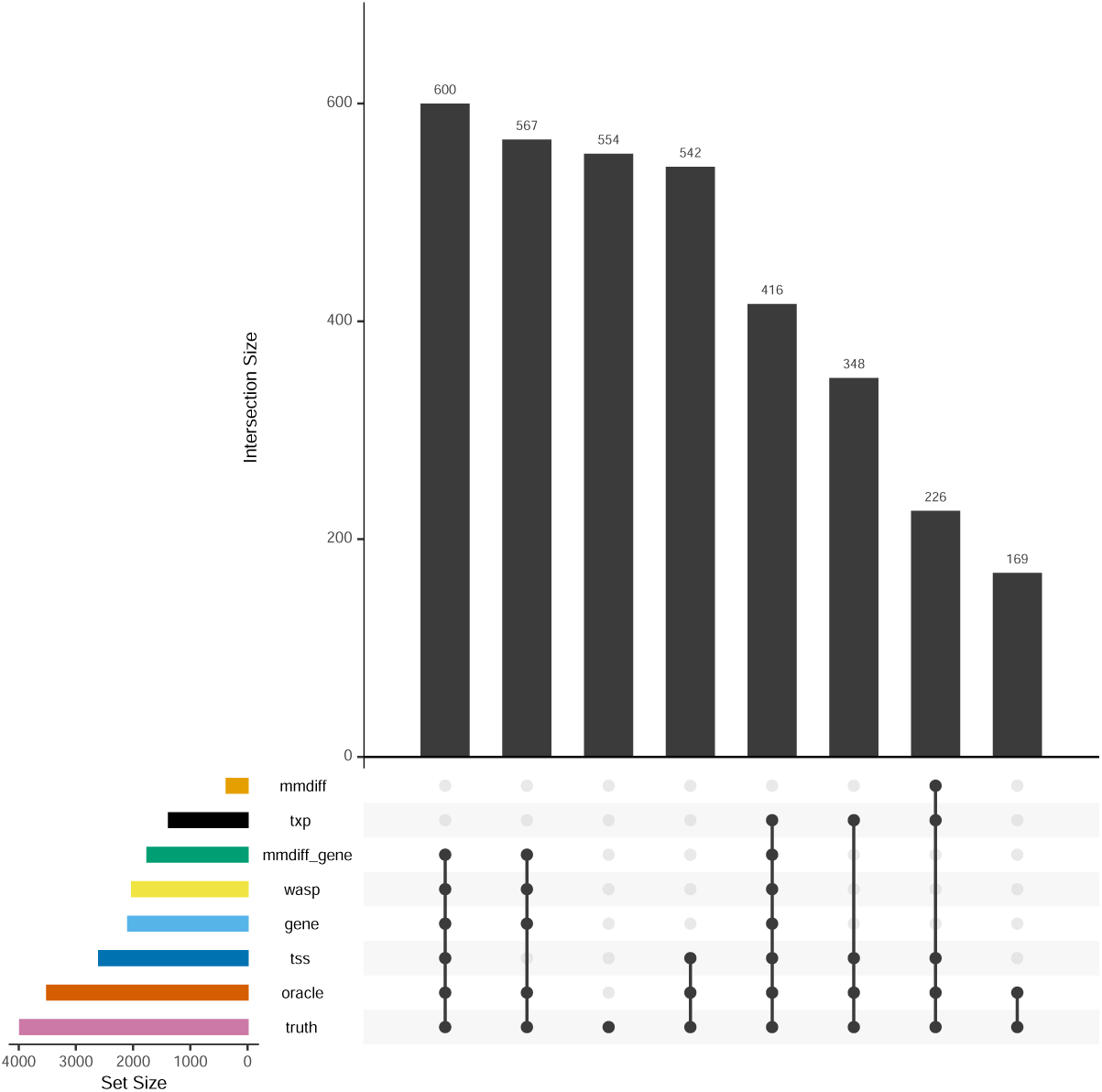
UpSet plot showing the overlap of simulation results at the transcript level comparing *SEESAW* at various levels of aggregation to *mmdiff* and *WASP*. The overlap is shown of different methods’ positive sets as well as the true AI status of each transcript.

**Figure S3.**
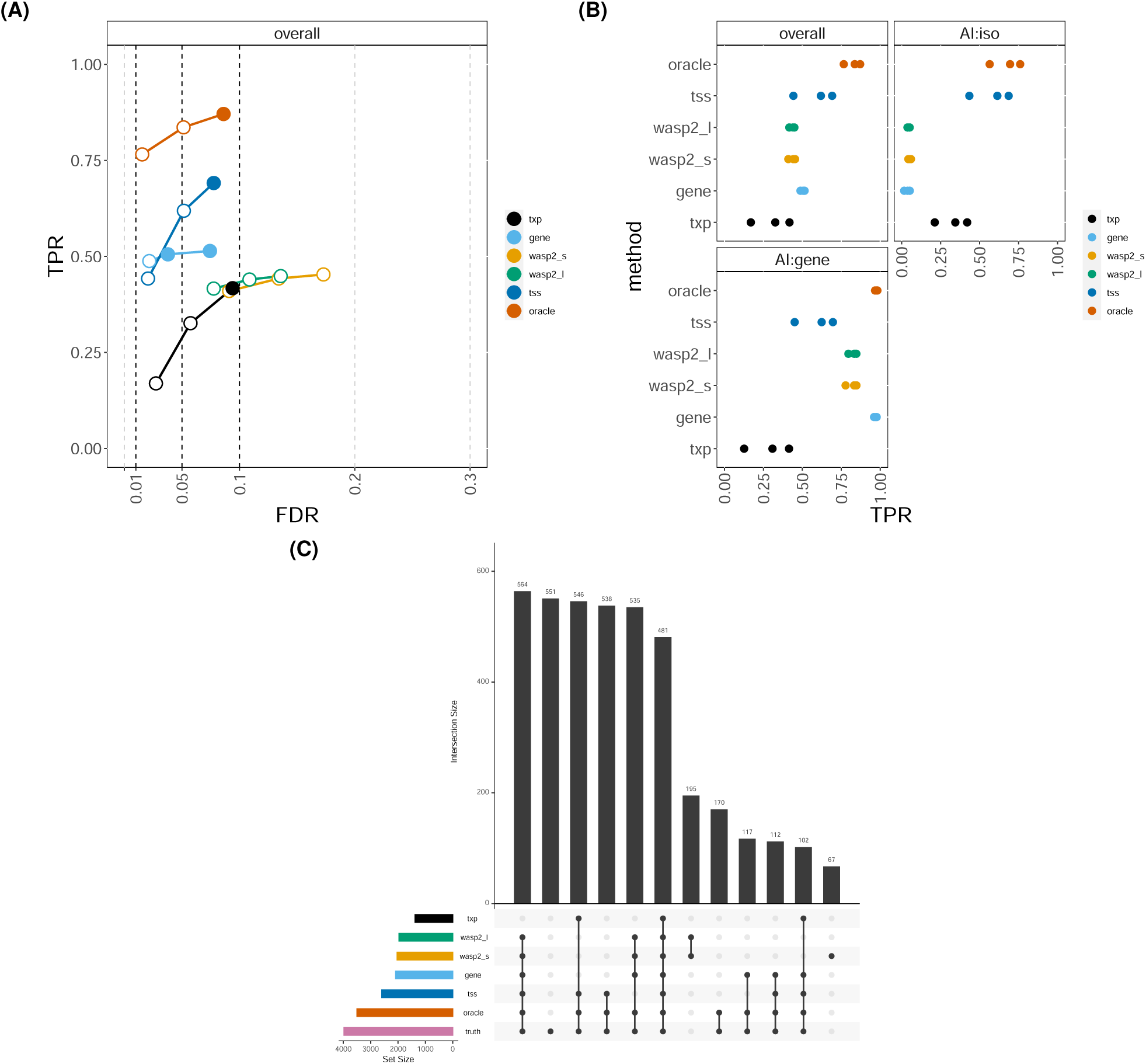
Simulation results at the transcript level comparing *SEESAW* at various levels of aggregation to *WASP2*. Both “single” (s) and “linear” (l) models in *WASP2* were assessed. A) Overall sensitivity over false discovery rate. B) Sensitivity overall and stratified by type of simulated AI. C) UpSet plot of the overlap of transcripts called by the methods and true AI transcripts.

**Figure S4.**
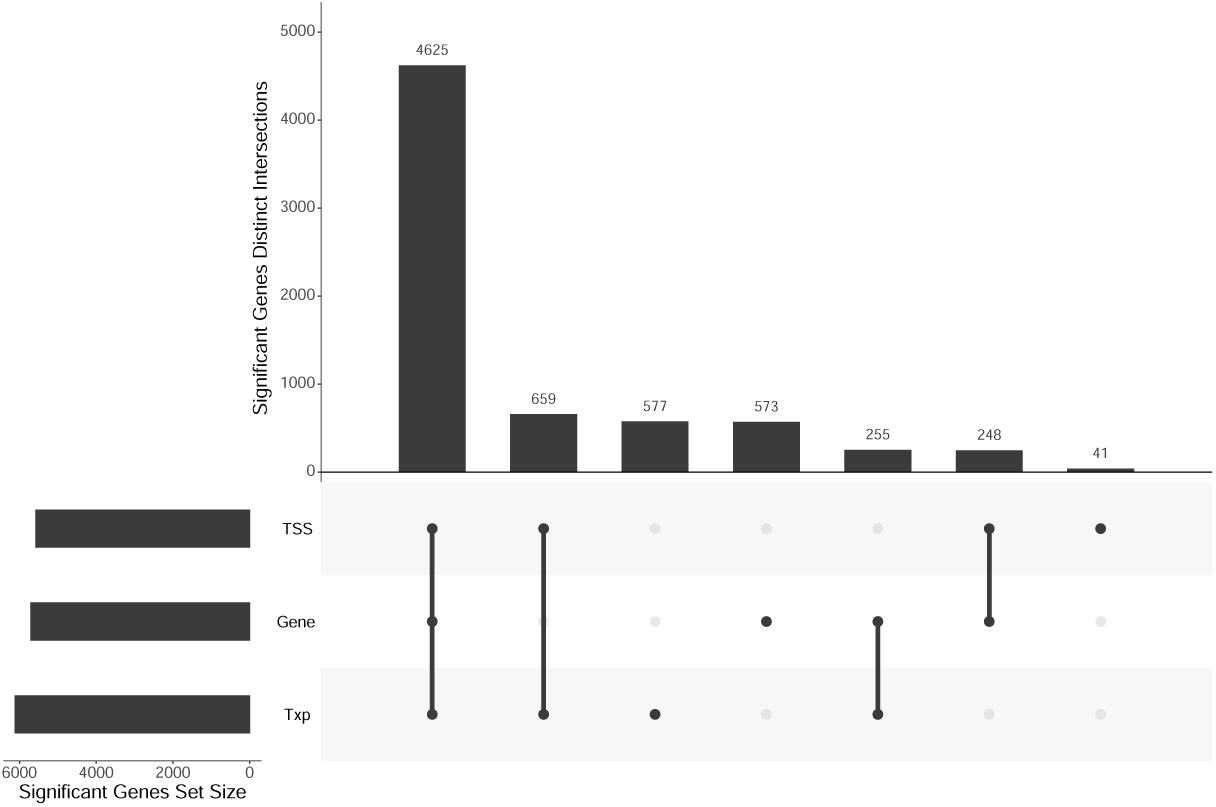
For the mouse osteoblast experiment, UpSet plot showing the overlap of significant genes, testing for global AI at various levels of aggregation using *SEESAW*.

**Figure S5.**
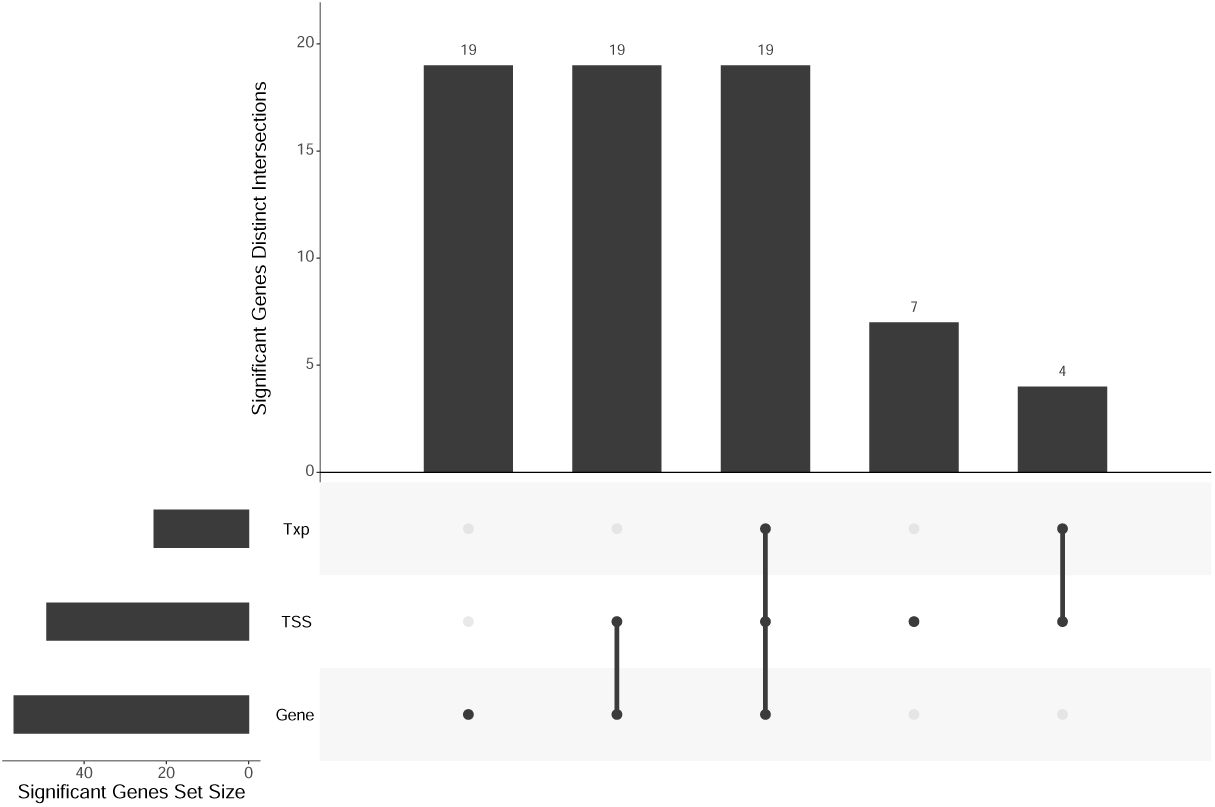
For the mouse osteoblast experiment, UpSet plot showing the overlap of significant genes, testing for dynamic AI at various levels of aggregation using *SEESAW*.

**Figure S6.**
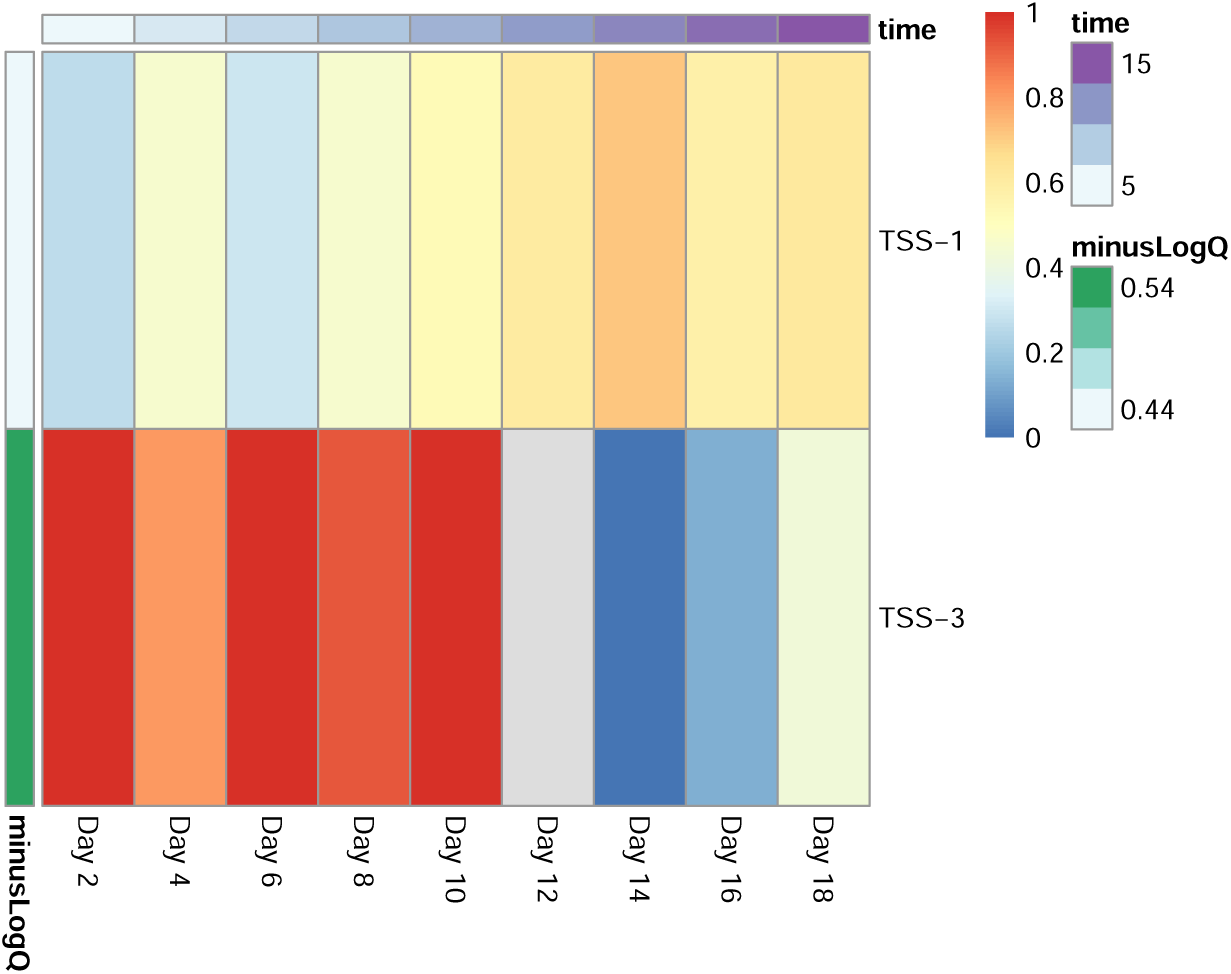
Allelic heatmap for two TSS groups of *Rasl11b*. minusLogQ denotes the *−* log_10_(*q*-value) for dynamic AI testing for each TSS group. Color indicates the fraction of total expression from the CAST/EiJ allele.

**Figure S7.**
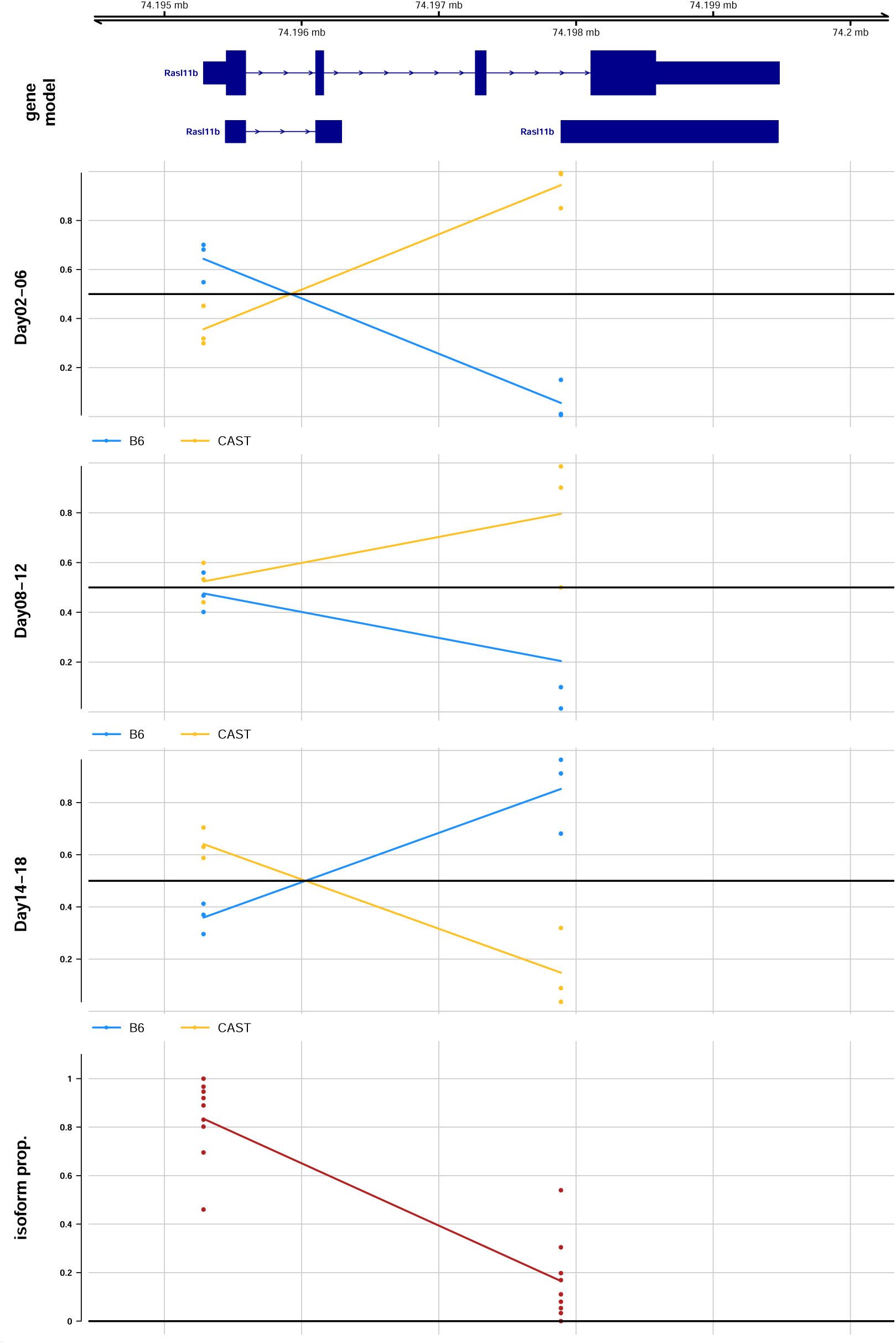
Gene model plot for *Rasl11b* tested at TSS-group level. Dynamic allelic ratios over three grouped time points: day 2-6 (top row), day 8-12 (middle row) and day 14-18 (bottom row). Isoform proportions from all days shown in the fourth row.

**Figure S8.**
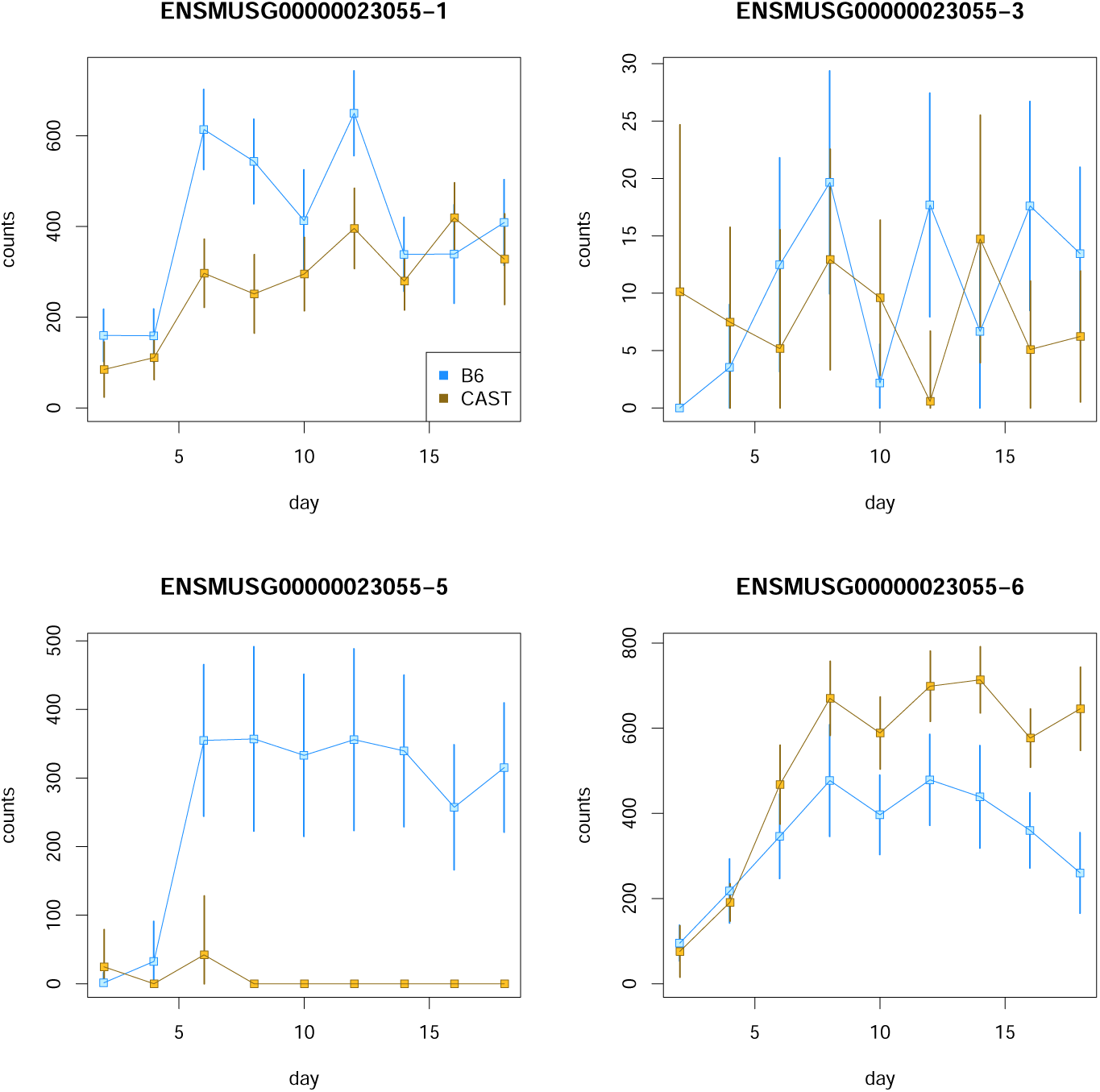
Estimated allelic counts over time for TSS groups of *Calcoco1*. Shown are four TSS groups of which “5” and “6” were significant for dynamic AI (FDR <5%). Estimation uncertainty shown with error bars (95% intervals based on bootstrap variance).

**Figure S9.**
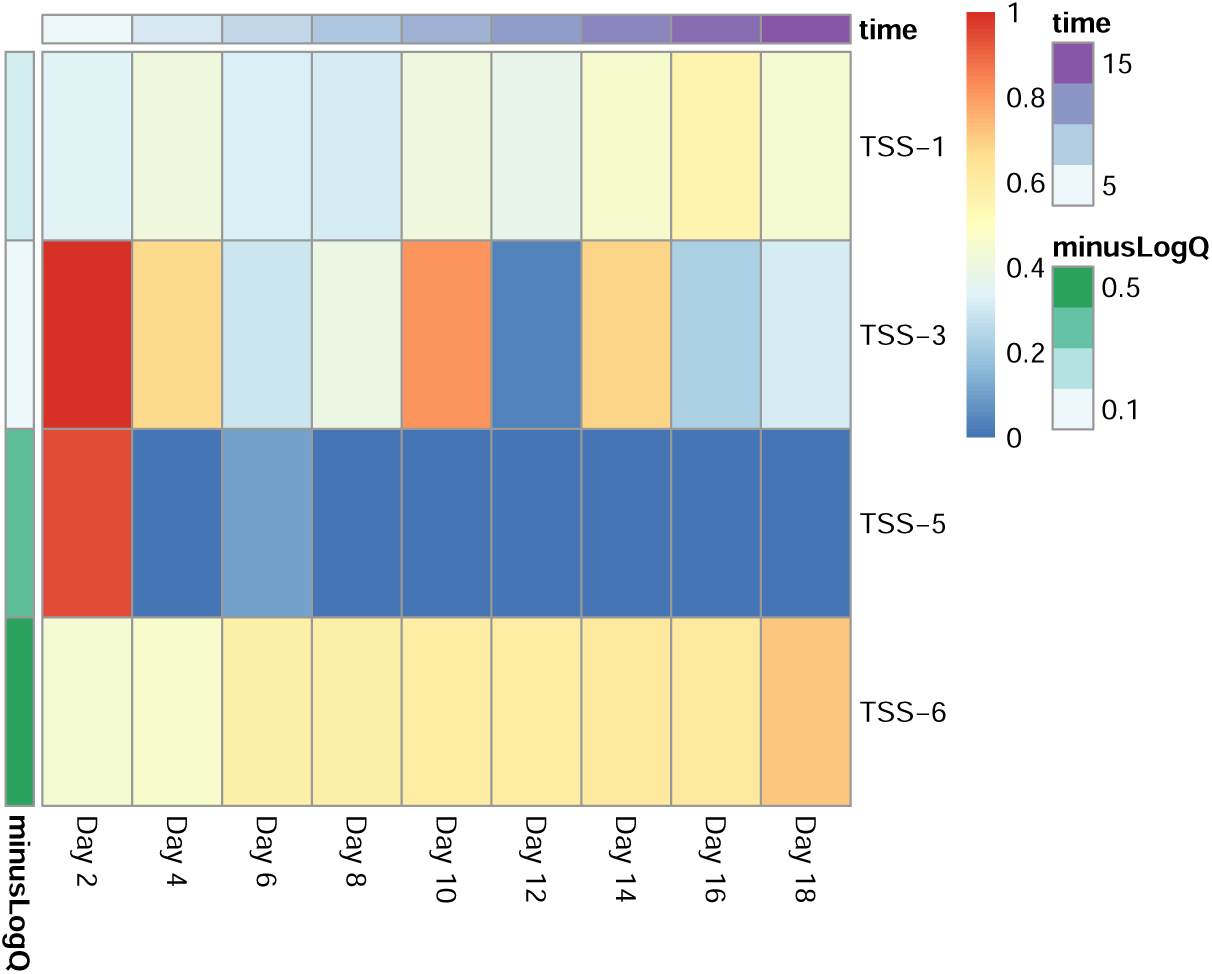
Allelic heatmap for four TSS groups of *Calcoco1*. minusLogQ denotes the *−* log_10_(*q*-value) for dynamic AI testing for each TSS group. Color indicates the fraction of total expression from the CAST/EiJ allele.

**Figure S10.**
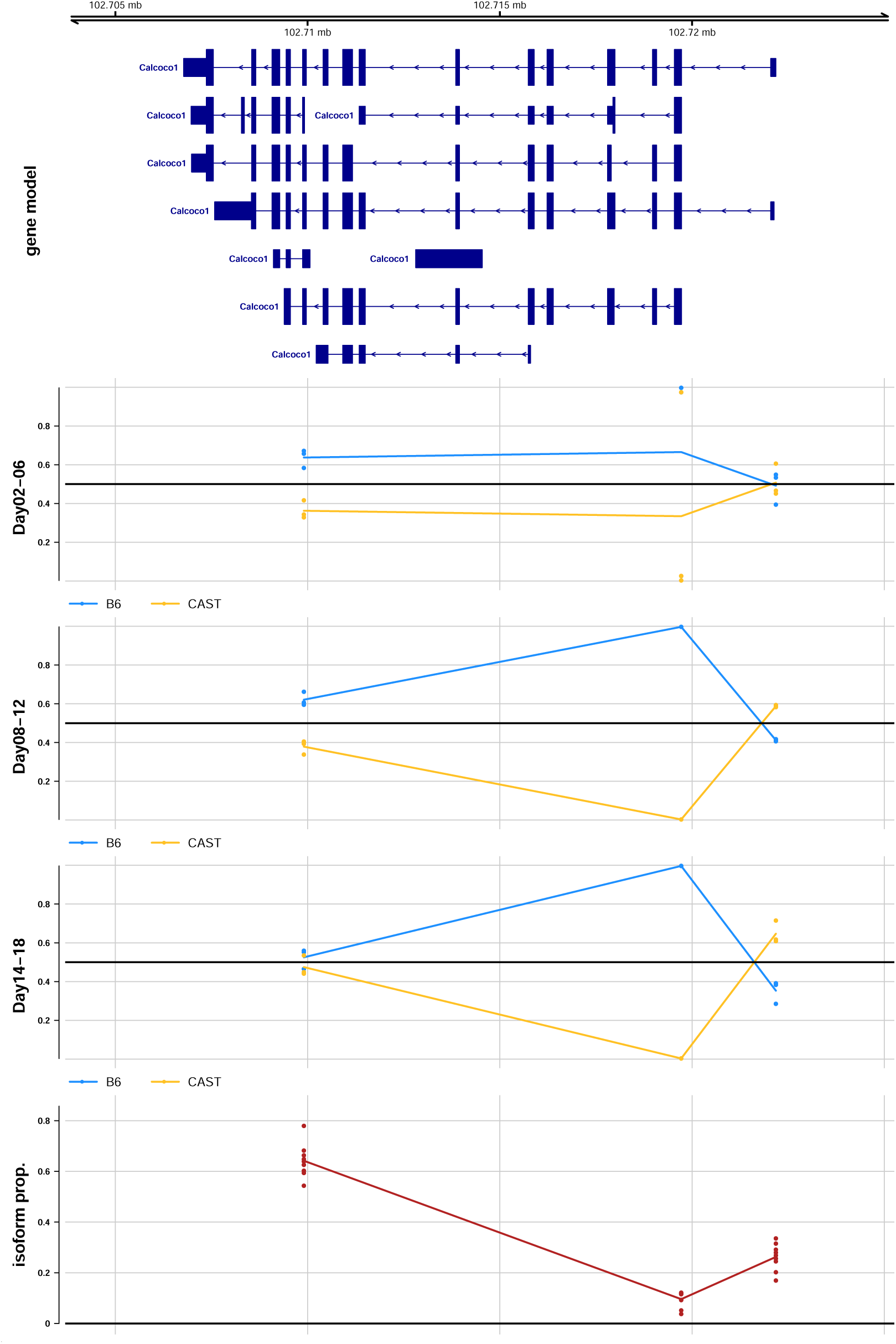
Gene model plot for *Calcoco1* tested at TSS-group level. Dynamic allelic ratios over three grouped time points: day 2-6 (top row), day 8-12 (middle row) and day 14-18 (bottom row). Isoform proportions from all days shown in the fourth row. From left to right, the TSS groups are “1”, “5”, and “6”. TSS group “3” was filtered in this plot due to too low counts and isoform proportion.

**Table S1.**
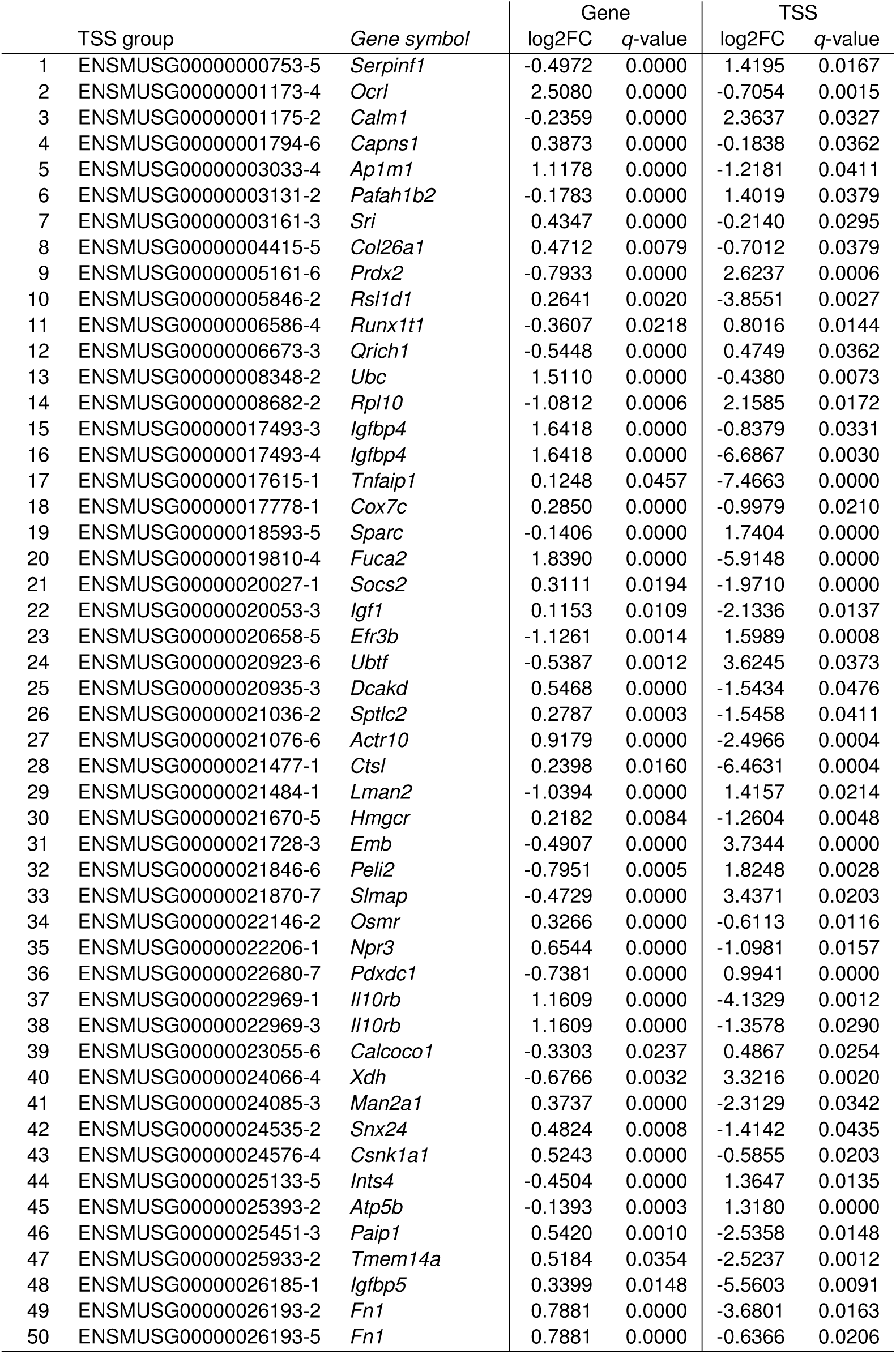
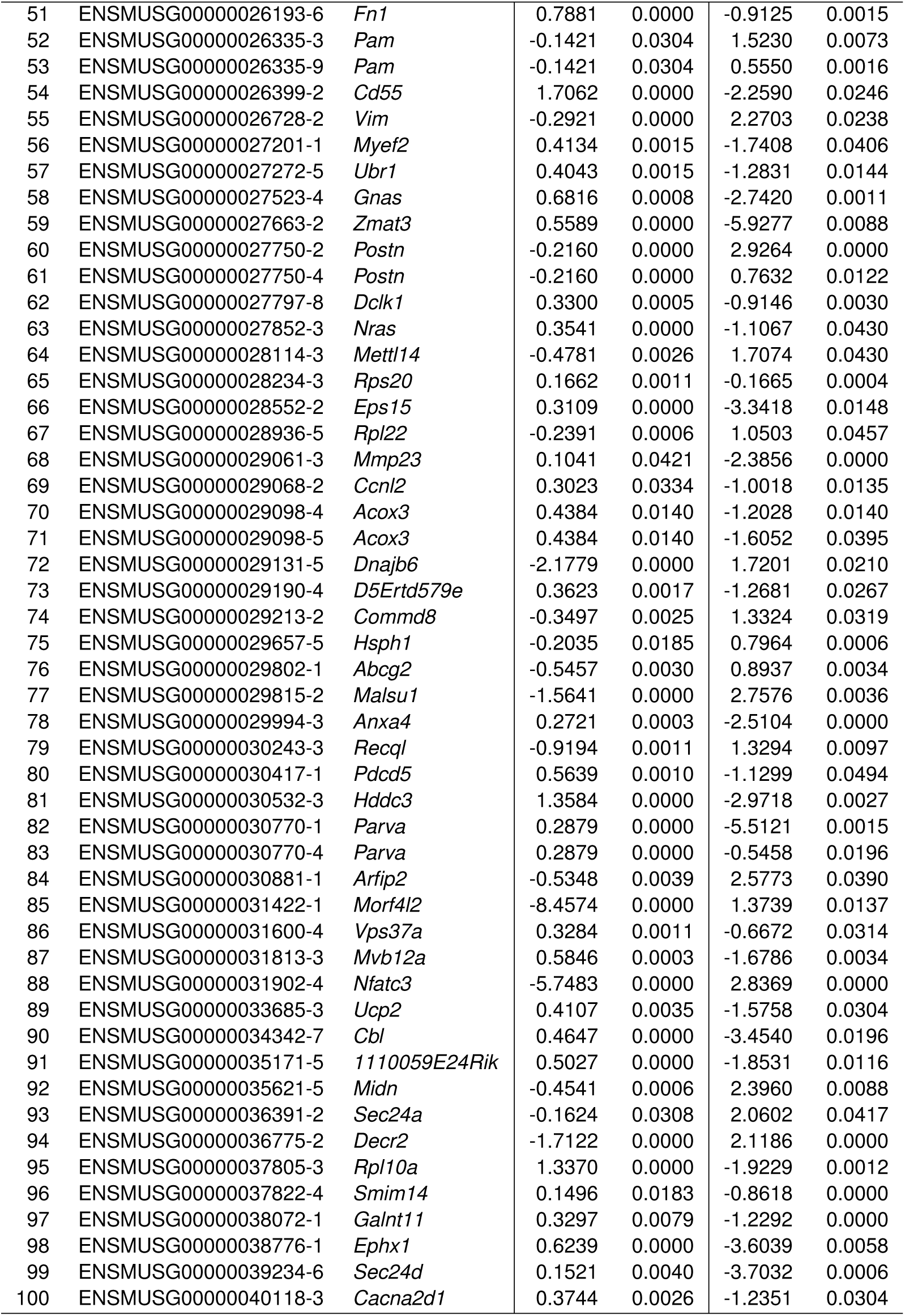
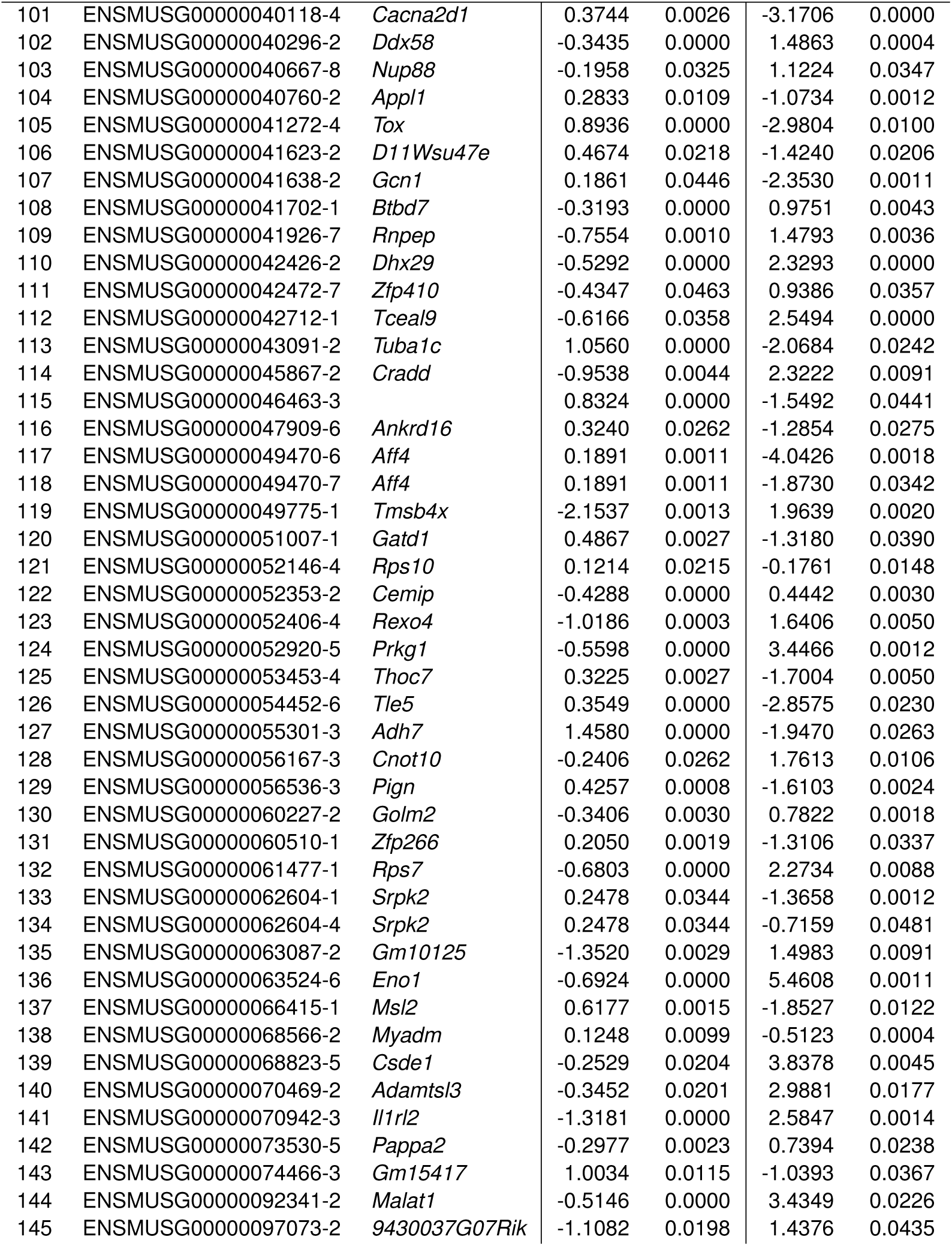
Genes that exhibited global AI with discordant direction of AI at gene and TSS-group level. Each row represents a TSS-group that was found significant for the global AI test (FDR <5%) and which had discordant sign compared to the gene-level AI (also FDR <5%).

